# Automation of high-throughput arrayed mammalian cell line cultivation

**DOI:** 10.1101/2025.10.03.676043

**Authors:** Chih-Cheng Yang, Aniruddha J. Deshpande, Michael Jackson, Peter D. Adams, Daniel Lynch, Amy V. Gibson, Zarina K. Waqar, Anna Beketova, Jiang-An Yin, Chun-Teng Huang

## Abstract

Cell culture automation has traditionally been limited to basic tasks at low throughput, which are insufficient for passaging rapidly proliferating cell lines or for generating stable clonal lines. To address unmet needs, this study implemented a Biomek i7 Hybrid automated workstation, integrated with peripheral instruments and coordinated by SAMI EX software, to enable automated, high throughput mammalian cell culture workflows. The workflows support cell density monitoring, arrayed passaging, sample cherry-picking, plate reformatting, cell density normalization, and cryopreservation in 96-well plates. Integration with the CloneSelect imager allows rapid confluency monitoring and monoclonality assessment (<100 sec per plate). Cell passaging and density normalization require 32 minutes for one plate and 61 minutes for two plates. Workflow consistency was demonstrated across multiple cell lines and biological replicates, with wells showing comparable confluency within three standard deviations, lower coefficient of variation, and substantially narrower interquartile ranges after a single cell passage and density normalization. Four automation pipelines, including monoclonality screening, cell passaging and cherry-picking, density normalization, and cryopreservation, collectively enable clonal line establishment. Depending on scale, one to eight 384-well plates were processed in 69 to 355 minutes, yielding an average of 35 clonal lines per plate suitable for downstream genomic DNA sequence confirmation.

## Introduction

The Biomek i-series automation model has been a trusted liquid handling platform since the mid-1980s and can be integrated with more than 300 devices from over 60 manufacturers to support different laboratory applications. The Biomek automation system has demonstrated its versatility in a wide range of workflows, including mammalian and stem cell culture maintenance [1, 2, 3, 4, 5], high throughput screening [3, 4, 6, 7, 8], assay processing [3, 9, 10, 11], sample treatment and reformatting [12, 13, 14, 15, 16], PCR and quantitative PCR [5, 7, 17, 18, 19], mRNA and ChIP-Seq library preparation [20, 21, 22], genomic and proteomic workflows [23, 24, 25, 26, 27, 28] and others.

In the past, automation using several Biomek series models enabled mammalian cell line cultivation in CELLSTAR AutoFlasks (Greiner Bio-One) for suspension cells [1], 3D cell cultures [4], and mouse embryonic stem cells in a 6-well plate format [2]. However, the throughput for these cell culture passaging processes was considered low. For higher throughput formats, such as 96-well plates, Biomek automation was primarily used for medium exchange to maintain induced pluripotent stem cells for downstream cell imaging or assay analysis, which did not require cell dissociation or passaging [3, 5]. In oncology studies, cancer cell lines typically exhibit rapid growth, with cell doubling times approximately 12 to 72 hours. As a result, cells reaching over-confluency became a major constraint in arrayed library screens using ORF (Open Reading Frame), shRNA (short hairpin RNA), and CRISPR-gRNA libraries. The effective screening window was limited to one week or less. This underscores unmet needs for high throughput automation of arrayed cell passaging and cell number normalization to extend the library screening timeframe. In addition to CRISPR library screening, CRISPR/Cas technology was widely adapted in biomedical research, particularly gene editing. Traditionally, screening and establishing stable clonal lines from CRISPR-mediated knockout or knock-in (KI) cell populations has been labor-intensive, time-consuming, and costly. To address these challenges, multiple high throughput pipelines were developed in this study utilizing the Biomek i7 workstation to automate cell confluency screening, arrayed cell passaging, sample cherry-picking, followed by plate reformatting and cell number normalization for cryopreservation. These pipelines enable the long-term maintenance and passaging of cell cultures as experimentally needed.

The arrayed mammalian cell culture automation workflow was established using the Beckman Coulter software suite, specifically Biomek 5 and SAMI EX software (v5.0), on the Biomek i7 Hybrid liquid handler platform. The Biomek 5 software was used to develop sophisticated liquid handling methods, whereas SAMI EX software can coordinate the Biomek workstation with peripheral instruments such as the washer, shaker, incubator, imager, and centrifuge. The scheduling interface in SAMI EX can manage and prioritize the hardware movement and method execution efficiently and in parallel. SAMI also supports method development and provides workflow simulation to test methods, identify errors, and troubleshoot before execution. This platform enabled fully automated, high throughput workflows optimized for arrayed mammalian cell line cultivation, described in this study, starting from the growth monitoring of single cell per well in a 384-well format to the cell panel expansion in a 96-well plate format.

## Materials and Methods

### 1. Hardware layout of the Biomek i7 Hybrid automated workstation and integrated instruments

The Biomek i7 Hybrid automated liquid handling workstation (Product No. B87585, Beckman Coulter) contains a hybrid multi-channel dual-pod system which combines 96/384-channel pipette heads and eight independent pipette heads, along with a barcode reader, two track grippers, a plate shuttle station, a Biotek 405 LSHTV washer (Agilent), a BioShake 3000-T ELM microplate shaker (QInstruments), a Cytomat 2C hotel incubator (Thermo Fisher Scientific), a CloneSelect cell imager (Molecular Devices), and a microcentrifuge (Agilent) (Figure 1A). These components work synergistically to streamline high throughput automation within a sterile enclosure with HEPA-filtered fans. The Beckman Coulter software suite includes Biomek software 5 for method and workflow development, as well as SAMI software for centralized workflow control, scheduling, and extended method design.

**Figure 1:**
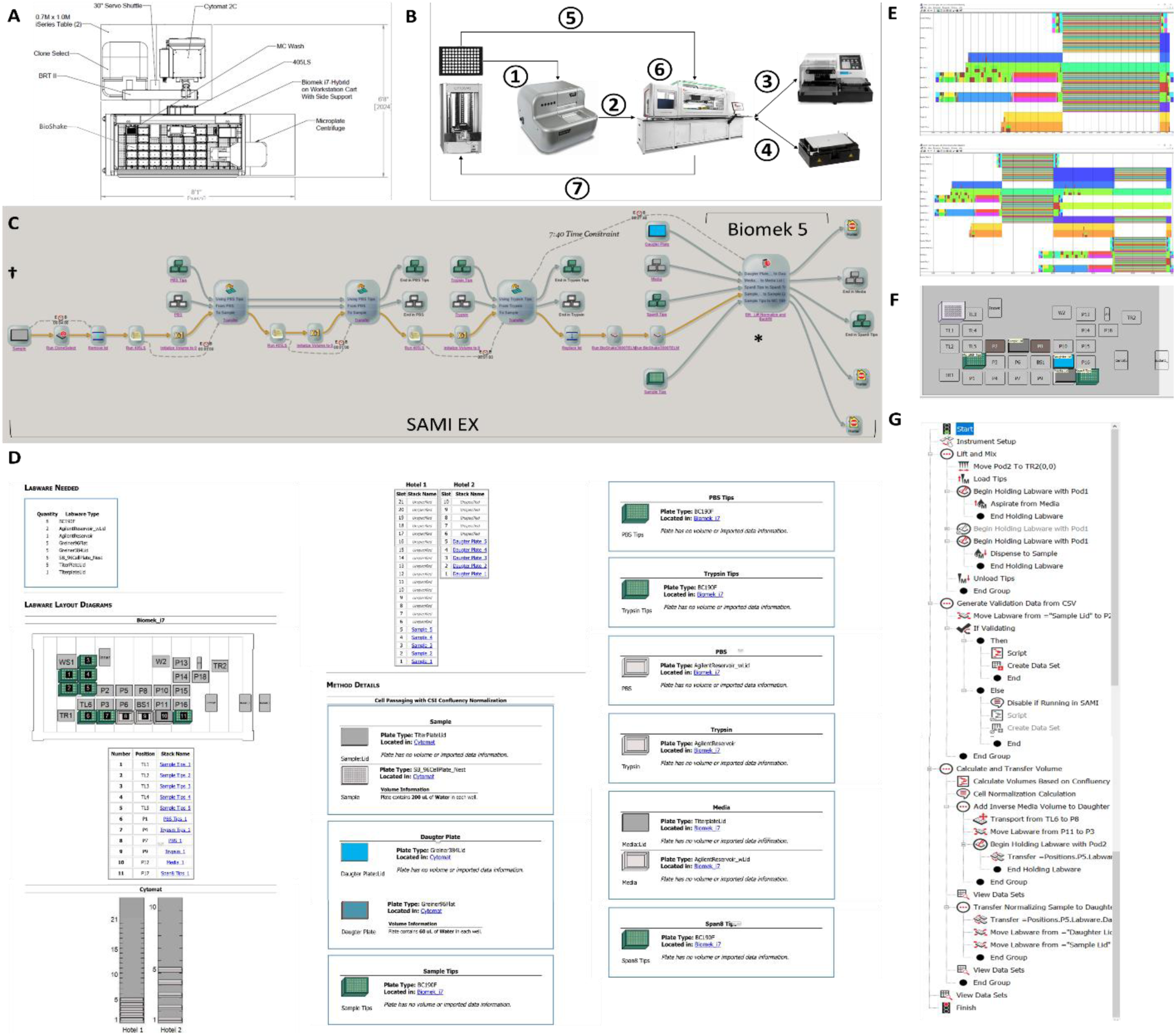
High-throughput automated arrayed cell culture passaging workflow. **A.** Hardware layout of the Biomek i7 Hybrid automated workstation. **B.** Schematic overview of arrayed cell culture passaging. ① A parental 96-well plate containing cells was transferred from the Cytomat 2C to the CSI via BRT II. ② The parental plate was transferred to the deck position of the Biomek i7 via the servo shuttle and track gripper. ③ The parental plate underwent two PBS washes using the 405LS washer and the Biomek i7 for dispensing PBS and trypsin. ④ Plate incubation at 37°C and vortexing during trypsinization were carried out using the BioShake. ⑤ A new daughter plate was transferred from the Cytomat 2C incubator to the Biomek i7 deck. ⑥ Media addition and mixing during the neutralization and backfill steps, as well as cell suspension transfer, were performed using the Biomek i7. ⑦ Both parental and daughter plates were transferred back to the incubator. **C.** Workflow overview under the SAMI EX interface. **D.** Overview of the SAMI EX deck layout, marked by the dagger symbol in Figure 1C. Notably, water was set as the default liquid type for all liquid reagents with comparable viscosity. **E.** Workflow scheduling and time estimation by SAMI EX illustrate the timestamps of each labware for cell passaging of a single (top panel) or two (bottom panel) 96-well plates in chronological order. **F.** The deck layout of the Biomek 5 method for cell density normalization and media backfill is shown at the position indicated by the asterisk symbol in Figure 1C. **G.** Overview of the Biomek 5 method for cell density normalization and media backfill, indicated by the asterisk symbol in Figure 1C.

**Figure 2:**
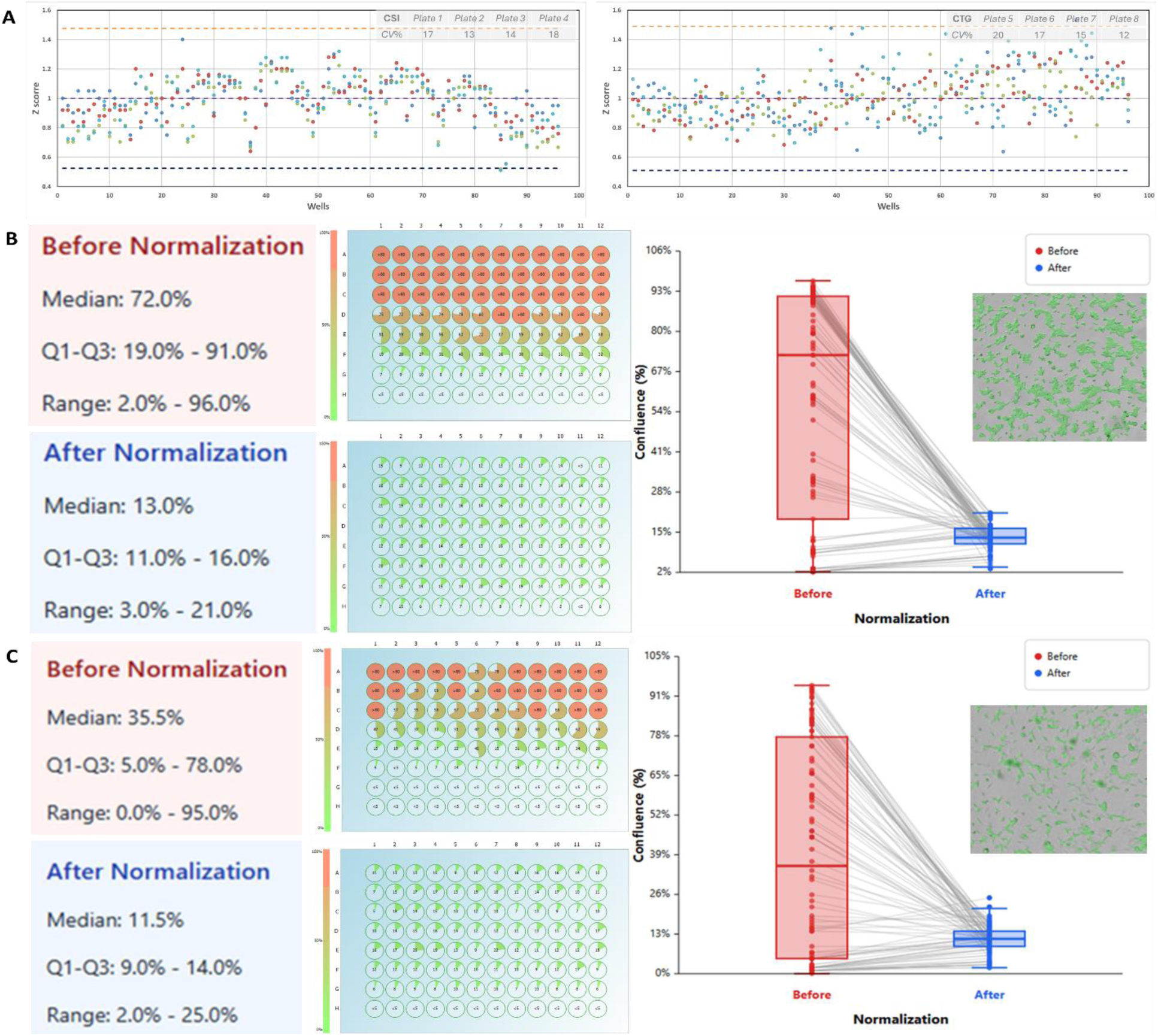
High-throughput arrayed cell density normalization. **A.** Comparison of CSI imaging and CTG viability assays for HEK293 cells in 96-well daughter plates with biological quadruplicates following cell density normalization of parental plates. The purple dashed line corresponds to the mean. The dashed orange and blue lines indicate three standard deviations above and below the mean, respectively. **B.** Comparison of HEK293 cell confluency before and after density normalization, assessed by CSI imaging and displayed as heatmaps. Each row, from top to bottom, represents a 2-fold serial dilution, with each column serving as a biological replicate (12 wells per dilution). Well confluency before and after normalization is also shown in a comparative boxplot. Representative cell image (right) from a well of the parental plate before normalization, with cell area masked in green and confluency analyzed using the CSI built-in algorithm. Parental plate incubated one extra day before normalization, compared with that in Figure 2C. **C.** Heatmap and boxplot comparison of HEK293T cell confluency before and after density normalization using CSI imaging. Rows represent 2-fold serial dilutions; columns are biological replicates (12 wells/dilution). Representative CSI image (right) from a well of the parental plate before normalization.

### 2. Arrayed mammalian cell culture passaging

The layouts of the incubator and the liquid-handling deck are shown in Figure 1D. This cell culture passaging workflow was set up for one or two 96-well plates per set and ran sequentially with limited overlapping steps. The number of plates may be increased but could be limited by the available deck and incubator space. A parental 96-well plate containing cultured cells ready for passaging was transferred from the incubator to the CloneSelect Imager (CSI) (Molecular Devices) using the Beckman Robotic Transport II (BRT II). The CSI measured cell confluency at high throughput (approximately 90 seconds per plate) and exported the cell density data as a CSV file to the Biomek i7 for passaging calculations. After imaging, the BRT transported the plate through a barcode scanner for hardware tracking and data recording. A liquid-handling gripper then placed the plate on the designated deck position, removed the lid, and transferred the plate to the 405 LSHTV Washer (BioTek) for media removal. After the plate was placed back on the deck, 150 µL of 1X PBS without magnesium and calcium was gently dispensed using a 96-channel pipette head at 100 µL/sec speed, with each tip touching the side wall of the well. The used tips were rinsed with sterile water via a MultiChannel wash station (MC Wash) before being ejected back into the tip box for future reuse. The plate was then transferred to the washer for PBS aspiration, and the entire wash procedure was repeated once. After the final PBS removal, 50 µL of 0.25% trypsin was added using a 96-channel pipette head with new tips, and the used tips were rinsed via the MC Wash. The lid was placed back on the plate, and the plate was incubated at 37 °C on a BioShake 3000-T ELM microplate shaker. After 5 min of incubation, cells were detached from the bottom of the well by vortexing on the shaker at 600 rpm for an additional 30 sec. After cell trypsinization, 150 µL of completed media was dispensed and mixed by pipetting up and down vigorously at 300 µL/sec speed for 100 times using the 96-channel pipette head, with the tip positioned 0.5 mm above the well bottom at 5 different coordinates within each well (20 times each in center, right, left, top, and bottom positions). In the meantime, a new daughter 96-well plate was transferred from the incubator, passed through the barcode reader, and placed on the designated deck position. The calculated volume of completed media was back-filled into each well of the daughter plate, followed by the transfer of a specific volume of cell suspension, each containing a normalized number of cells, from the wells of parental plate to their corresponding wells of the daughter plate using a flexible 8-channel pipette. Both parental and daughter plates were transferred from the deck position, passed through the barcode reader, and returned to the incubator (Figure 1B).

#### 2.1. Calculation of suspension volumes for mammalian cell passaging

Cell confluency data from each CSI imaging session were recorded as numerical percentages, exported as CSV files, and automatically transferred to the Biomek i7 system. Confluency values below 10% were adjusted to 10%. This lower-limit capping was implemented to limit the cell passaging ratio within a practical range of 1:1 to 1:10 and to avoid calculation errors with extreme values, such as 0% confluency or pipetting volume below 1 µL, which can occur during normalization between fully confluent (100%) and nearly empty (0%) wells. Confluency was then represented on a simplified 10-100 scale, with 100 denoting full (100%) well coverage. The Biomek 5 algorithm then applied

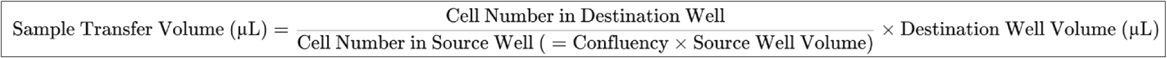

the formula to calculate the sample transfer volume from the parental (source) well to the daughter (destination) well, with each parameter user-configurable as either a variable or a fixed value. First, an arbitrary estimate of the total cell number per well in the parental plate was derived by multiplying the measured confluency by the respective source well volume (in µL). In this study, both the source and destination well volumes were 200 µL. When a source well’s confluency was 100 (representing 20,000 cells), targeting 2,000 cells in the destination well, which corresponds to a 1 to 10 cell passaging ratio, required a sample transfer volume of 20 µL, with a media backfill volume of 180 µL. On the other hand, when a source well’s confluency was at the minimum value of 10, the entire 200 µL cell suspension was transferred to the destination well.

### 3. Arrayed mammalian clonal line establishment

The high-throughput arrayed clonal line establishment workflow consists of four pipelines: 1) cell monoclonality screening, 2) clonal line passaging and cherry-picking, 3) cell number normalization for assay analysis and clone validation, and 4) clonal line cryopreservation.

#### 3.1. Arrayed mammalian cell monoclonality screening

The layouts of the incubator and the liquid handling deck are shown in Figure 3D. The maximum capacity for this workflow is two 384-well plates per set. After 12 hours of incubation, a set of two parental 384-well plates containing single-cell seeded cultures, originating either from limiting dilution (0.5 cells per well) or flow cytometry sorting (1 cell per well), was transferred from the incubator to the CSI using the BRT II for cell imaging at a predefined focal plane setting. Subsequently, the plate was imaged either daily or every several days, depending on the doubling time of the cells, to monitor cell growth (Figure 3A).

**Figure 3:**
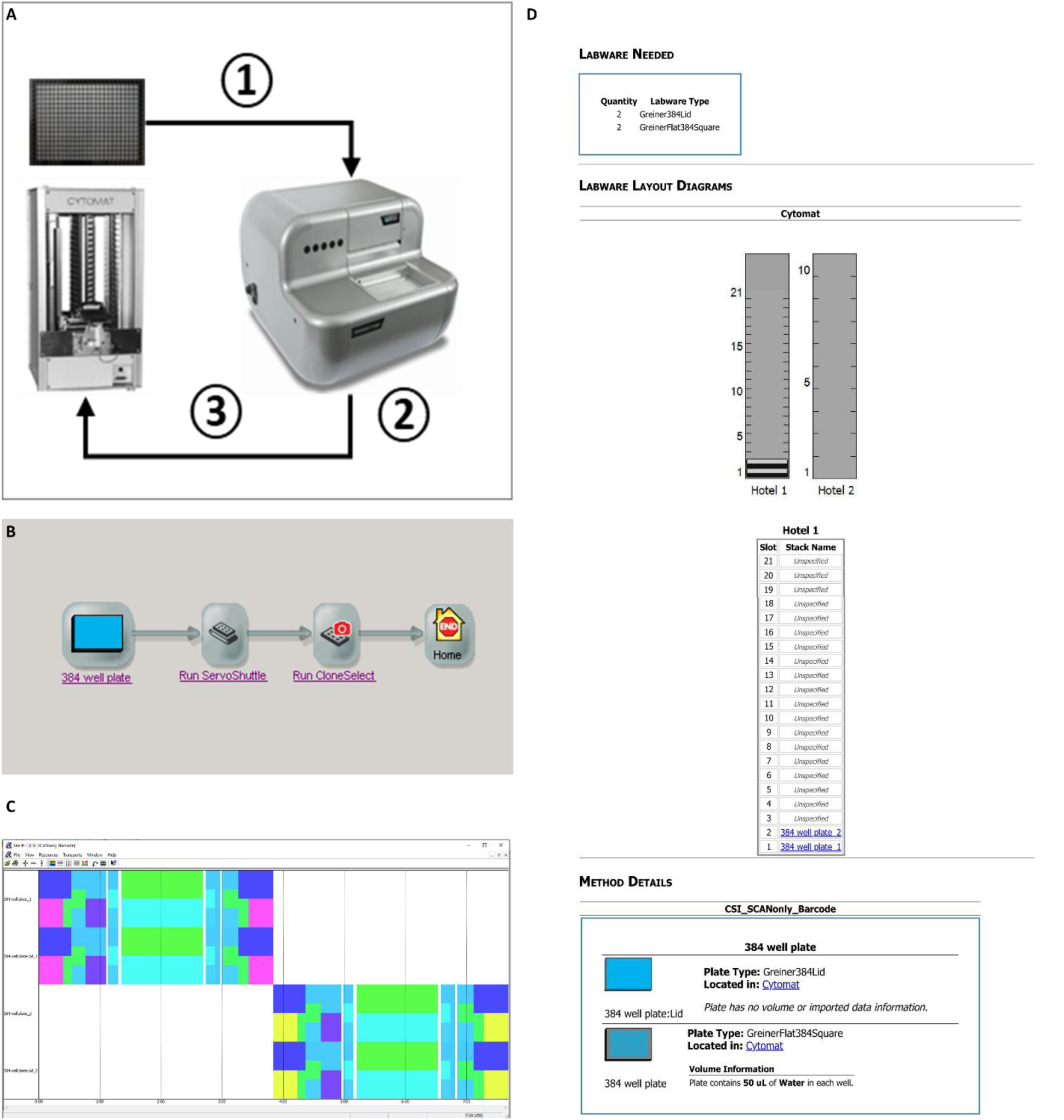
High-throughput automated arrayed cell monoclonality screening pipeline. **A.** Schematic overview of arrayed clonal line screening. ① A parental 384-well plate, seeded with a single cell per well, was transferred from the Cytomat 2C incubator to the CloneSelect imager via BRT II. ② High-throughput cell imaging via CSI. ③ After imaging, the plate was returned to the Cytomat 2C. **B.** Pipeline overview under the SAMI EX interface. The SAMI EX software was utilized for both method creation and workflow design. **C.** Pipeline scheduling, labware timestamps, and time estimation for clonal line screening of two 384-well plates were generated using SAMI EX. **D.** Overview of the SAMI EX labware layout for cell monoclonality screening.

**Figure 4:**
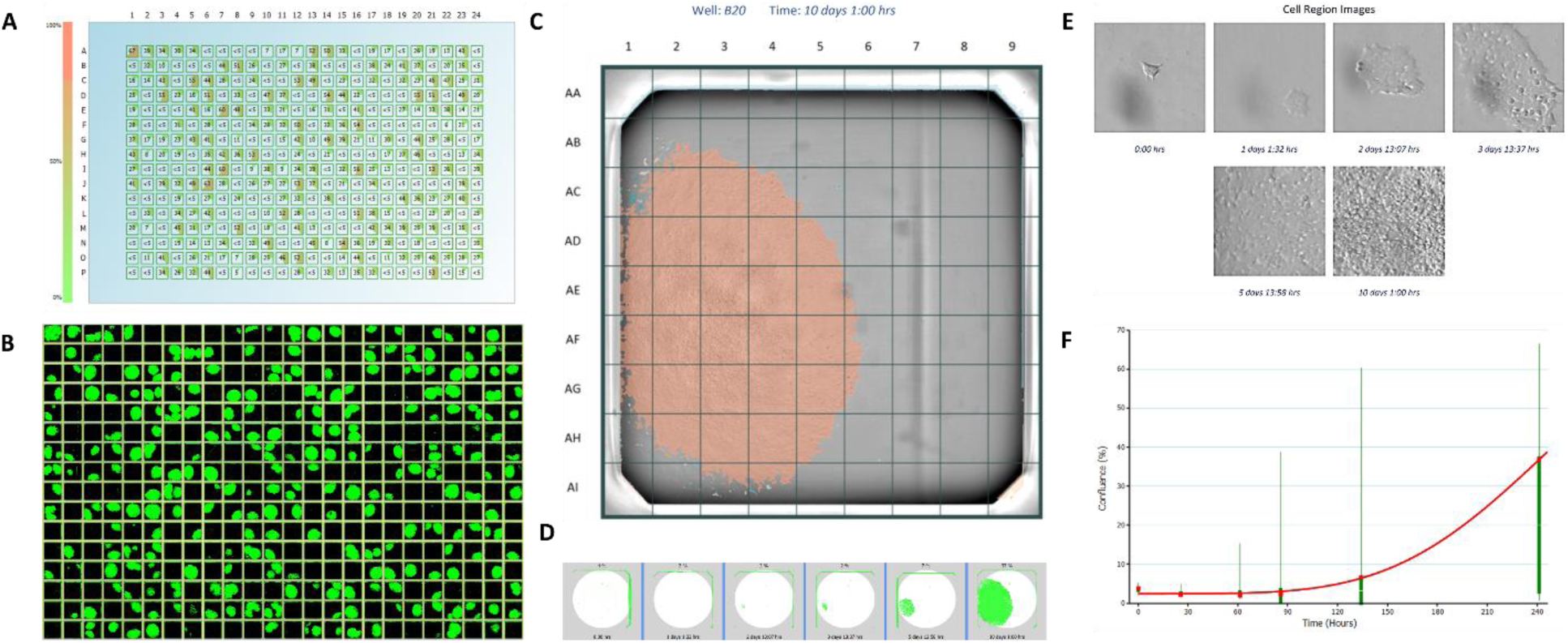
Monoclonality assessment for clonal line establishment. **A.** Cell confluency data, measured by CSI and its algorism, is presented both numerically and as a heat map. **B.** Cell confluency images are presented in thumbnail format. One hundred cells were seeded in the A1 position as the cell-focusing control well, while the remaining 383 wells were seeded with single cells. **C.** The cell confluency image for well B20 was captured 10 days post-seeding. A fixed total of 81 fields per well is available for cell tracking over time using time stamped images. **D.** Well confluency images for well B20 were time stamped and captured from day 0 (cell seeding) through day 10 post-seeding. **E.** Images of single-cell expansion in well B20 were time stamped and captured within a user-defined field from day 0 to day 10 post-seeding. **F.** Clonal growth curve of well B20 based on confluency from day 0 to day 10 post-seeding.

#### 3.2. Arrayed mammalian clonal line passaging and cherry-picking

The layouts of the incubator and the liquid-handling deck are shown in Figure 5D. Upon completion of the final imaging time point, approximately 1 to 2 weeks after cell seeding, cell confluency data was exported as a CSV file, along with the monoclonality report, to prioritize sample orders and generate a well-by-well cherry-picking report in an Excel spreadsheet containing true/false binary decisions for each sample well. Each parental 384-well plate underwent steps similar to the arrayed cell culture passaging workflow described in Step 2, including two consecutive 50 µL PBS washes (without a washer), 40 µL of trypsinization, 40 µL of media neutralization, and 35 µL of suspension mixing. All these steps were performed using a 384-channel pipette head. However, the pipeline was terminated in the neutralization step after mixing vigorously at 300 µL/sec speed for 100 cycles, with the tip positioned 0.1 mm above the well bottom at 5 different coordinates within each well (20 times each in center, and four corner positions). Instead of proceeding with the normalization procedure, wells containing cell suspensions of fast-growing, single-cell derived colonies flagged as “true” in the cherry-picking report were transferred at user-defined volumes and reformatted into a daughter 96-well plate format for clonal expansion (Figure 5A). The daughter plate was pre-filled with complete media in each well.

**Figure 5:**
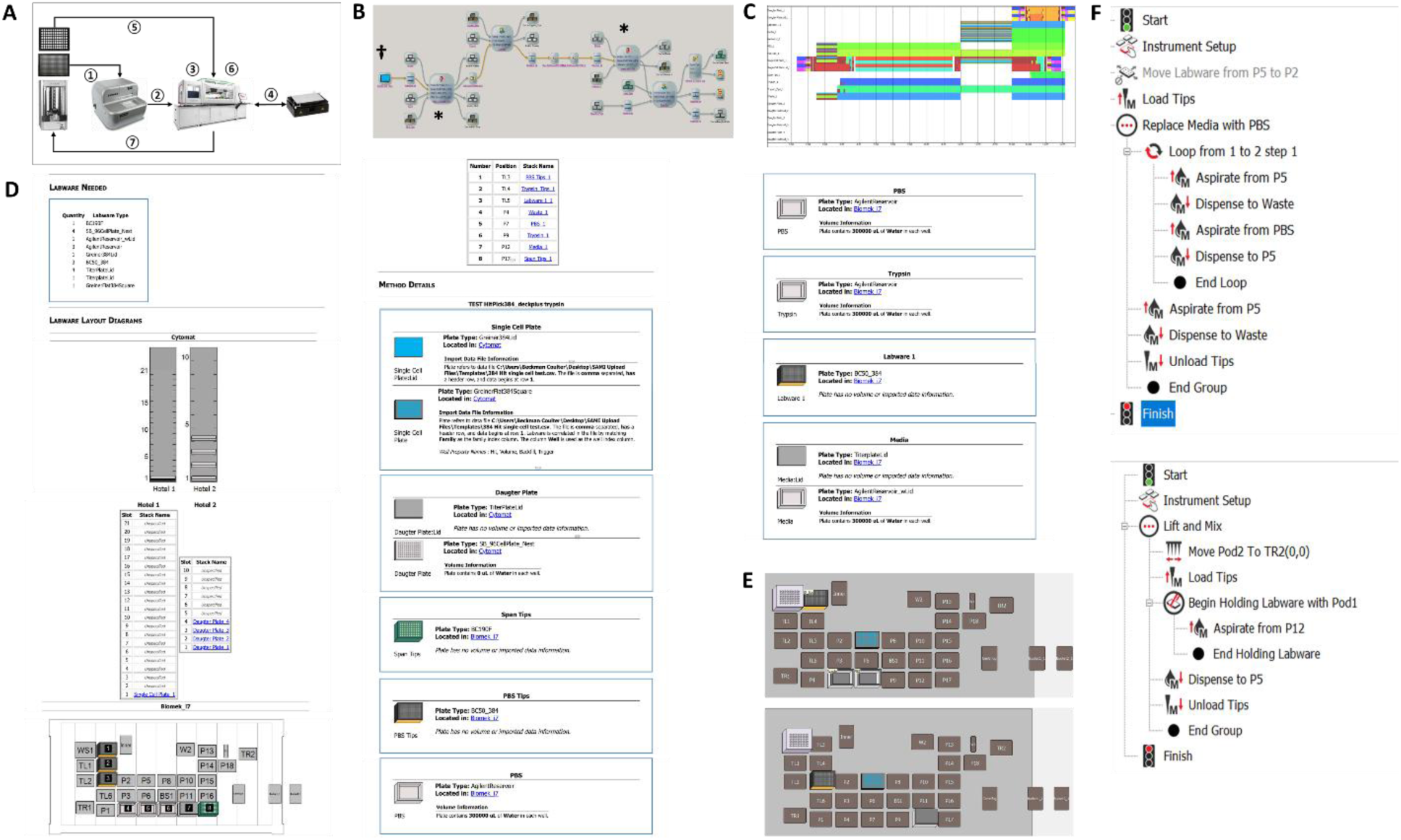
High-throughput automated arrayed clonal line passaging and cherry-picking pipeline. **A.** Schematic overview. ① A parental 384-well plate containing clonal line candidates was transferred from the Cytomat 2C incubator to the CloneSelect imager for confluency imaging via BRT II. ② The parental plate was transferred to the deck position of the Biomek i7 via the servo shuttle and track gripper. ③ The parental plate underwent two PBS washes, using the 384-channel pipette head for liquid removal and the same head for dispensing PBS and trypsin. ④ Plate incubation at 37°C and vortexing during the trypsinization step were carried out using the BioShake. ⑤ A new daughter 96-well plate was transferred from the Cytomat 2C incubator to the Biomek i7 deck. ⑥ Media addition and mixing during the neutralization and backfill steps, as well as well cherry-picking for cell suspension transfer, were performed using the Biomek i7. ⑦ Both parental and daughter plates were transferred back to the Cytomat 2C. **B.** Pipeline overview under the SAMI EX interface. Both SAMI EX and Biomek 5 software were utilized for the method creation and pipeline design. **C.** Scheduling and time estimation for the entire pipeline by SAMI EX illustrate the timestamps of each labware item in chronological order. **D.** The labware setup report, marked by the dagger symbol in Figure 5B, serves as a starting reference, providing a comprehensive overview of the SAMI EX deck layout, along with details on labware and associated conditions. **E.** The deck layout of the automated Biomek 5 methods specific to PBS replacement (top panel) and cell suspension mixing (bottom panel) are shown at the position indicated by the asterisk symbols in Figure 5B. **F.** Overview of the automated Biomek 5 methods for PBS replacement (top panel) and cell suspension mixing (bottom panel), indicated by the asterisk symbol in Figure 5B.

#### 3.3. Arrayed mammalian clonal cell number normalization for assay analysis and clone validation

The layouts of the incubator and the liquid-handling deck are shown in Figure 1D. The daughter 96-well plate then served as a new source plate and underwent the confluency screening pipeline (Step 3.1) to monitor cell growth. Once confluency was reached, the plate proceeded through the entire arrayed cell culture passaging workflow (Figure 1B), described in Step 2, to generate a new normalized 96-well plate (the granddaughter plate) for clone validation via assay analysis. The assays utilized in this pipeline were the CellTiter-Fluor (CTF) cell viability assay and the HiBiT assay.

#### 3.4. Arrayed mammalian clonal line cryopreservation

The layouts of the incubator and the liquid-handling deck are shown in Figure 7D. During clone analysis to identify suitable candidates using the granddaughter plate, 75 µL of fresh media were dispensed into each well of the passaged daughter plate containing leftover cell cultures from Step 3.3 using a 384-channel pipette head. After mixing by pipetting up and down, 75 µL of the resulting cell suspension was transferred from the passaged daughter plate to a new 96-well plate containing 75 µL of freezing agent (80% of FBS and 20% of DMSO) for cryopreservation and long-term storage (Figure 7A).

**Figure 6:**
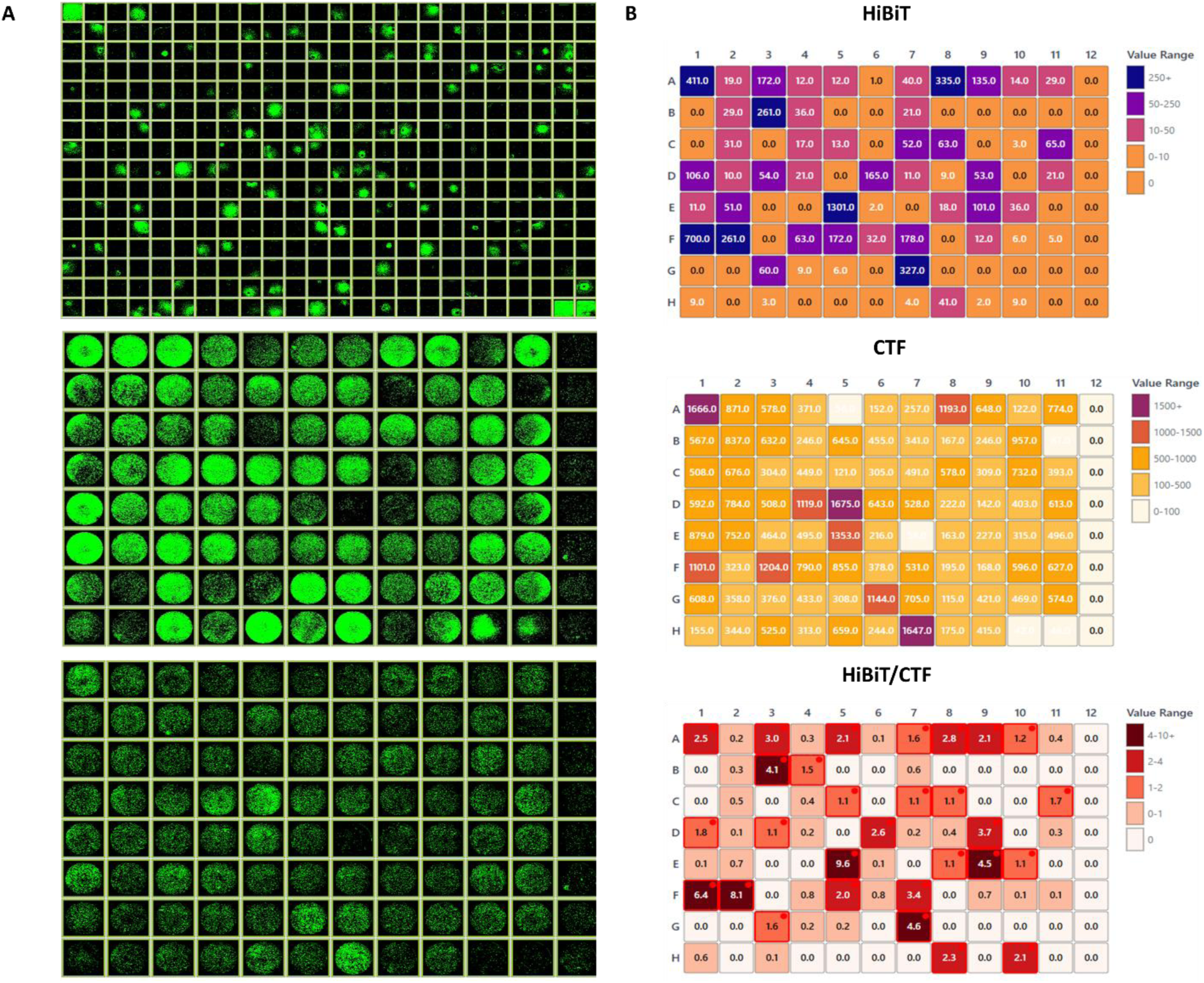
Arrayed mammalian clonal line establishment for CRISPR HiBiT tag knock-in. **A.** A series of cell confluency images shown in a thumbnail format from the monoclonality screening pipeline. The top image was captured 26 days post cell seeding in a 384-well plate and prior to passaging into a 96-well plate. The top image was acquired after 12 days of incubation following arrayed passaging and cherry-picking of mammalian clonal lines. The bottom image was captured after 1 days of incubation following arrayed mammalian clonal cell number normalization. **B.** CTF and HiBit assay analysis for the bottom 96-well plate in Figure 6A. Arbitrary values and Heatmaps displaying HiBiT signals (top figure) in relative luminescence units (RLU), CTF intensities (middle figure) in relative fluorescence units (RFU), and HiBiT normalized to CTF values (bottom figure). Wells with high normalized HiBiT values over 1 were selected as hits and labeled with red outline and a dot at the well corner.

**Figure 7:**
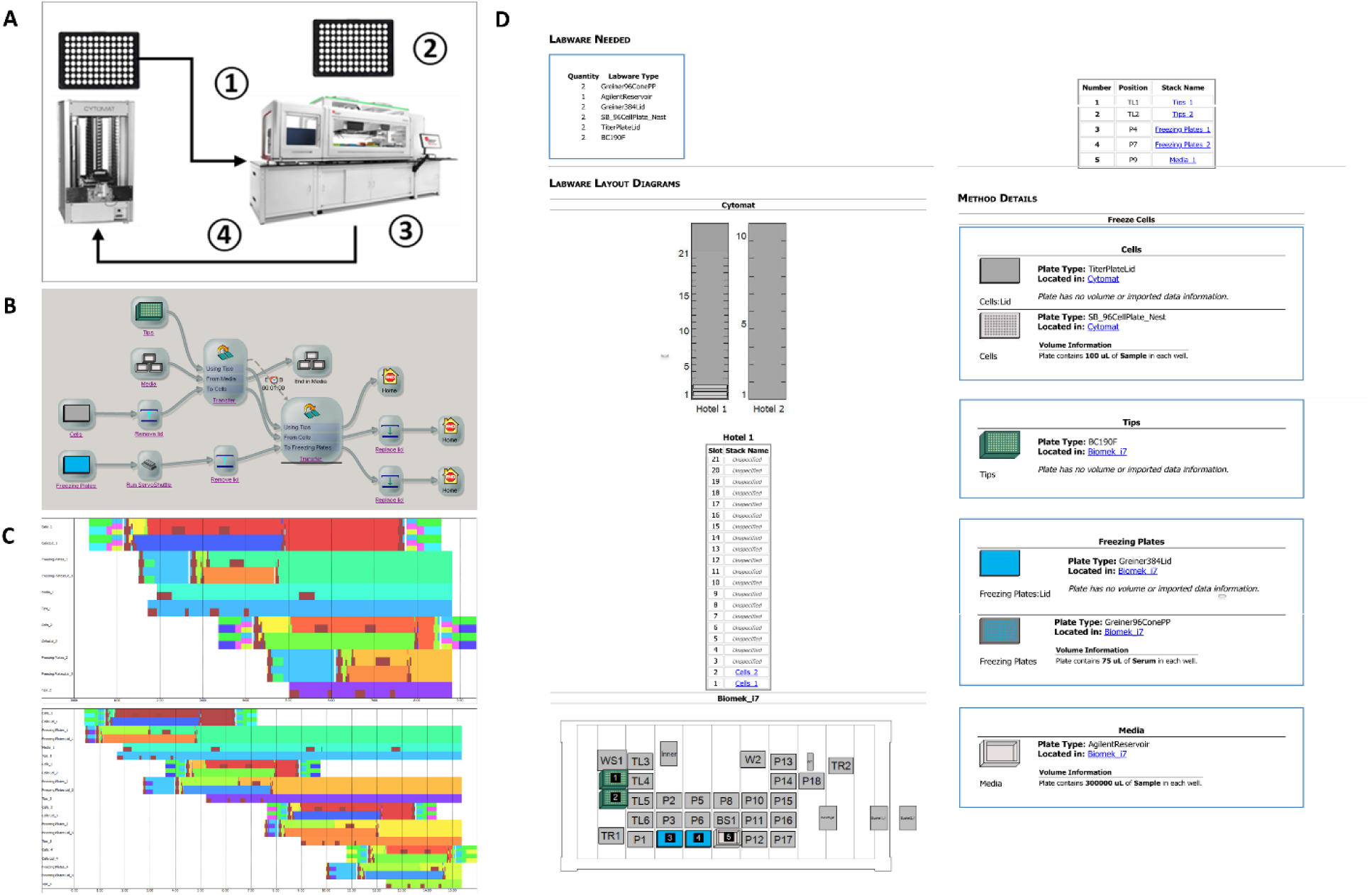
High-throughput automated arrayed clonal line cryopreservation pipeline. **A.** Schematic overview. ① The leftover passaged daughter 96-well plate from Step 3.3, which contained clonal cell suspensions, was transferred from the Cytomat 2C incubator to the CloneSelect Imager for confluency imaging using BRT II. ② A new granddaughter 96-well plate was positioned on the deck of the Biomek i7 as its home location. ③ Media, FBS, and DMSO addition and mixing, as well as well cell suspension transfer, were performed using the Biomek i7. ④ The daughter plate was transferred back to the Cytomat 2C, while the granddaughter plate remained in its deck position. **B.** Pipeline overview under the SAMI EX interface. The SAMI EX software was utilized for both method creation and workflow design. **C.** SAMI EX was used to generate pipeline schedules, labware timestamps, and time estimates for clonal line cryopreservation of one (top figure) and four (bottom figure) 96-well plates. **D.** SAMI EX labware layout designed for the cryopreservation pipeline of one 96-well clonal plate.

## Result

### High-throughput automation design

The prototype of the Biomek i7 Hybrid automated workstation is a dual-pod liquid handler. The right pod incorporates eight independent pipette heads to enable sample cherry-picking, media backfilling, and plate reformatting tasks in both cell normalization (Figure 1) and clonal line selection (Figure 3) workflows. The left pod is equipped with an interchangeable 96 or 384-channel pipette head to allow PBS, trypsin, media, and buffer dispensing, as well as high speed intra-well mixing to effectively disperse cell clumps during neutralization (Figures 1, 5, and 7). To enable automated, high throughput arrayed mammalian cell culture, this study integrated multiple instruments into the Biomek i7, and each equipment served a distinct function and purpose within the overall automation workflow design (Figures 1, 3, 5, and 7). The Cytomat 2C incubator accommodates two types of rack spaces (Figures 1D, 3D, 5D, and 7D), one for thinner plates such as 96– and 384-well plates, and the other for thicker plates ranging from 48-well to single-well OmniTrays. The CloneSelect imager (CSI) is designated for high throughput imaging of cell confluency and monoclonality assessment (Figure 4). Media and PBS aspirations are carried out using the BioTek 405 LSHTV washer, which enables rapid and thorough liquid removal. Trypsinization is facilitated by the BioShake 3000-T ELM, which provides both heating and orbital shaking to promote efficient detachment of the cell monolayer (Figure 1B).

### Arrayed mammalian cell culture workflow

The layout of the Biomek i7 system and integrated instruments is shown in Figure 1A. The workflow of arrayed cell passaging in a 96-well plate format is illustrated in Figure 1B, which can be easily optimized for different cell lines. Both Biomek 5 and SAMI EX software were used to develop methods and assemble a unified workflow under the SAMI EX interface (Figure 1C). Time constraints, indicated by dotted lines with time and clock icons, were embedded within or between methods to ensure uninterrupted workflow. The labware setup report (Figure 1D), indicated by the dagger symbol in Figure 1C, provided an initial overview of the SAMI deck layout, including the name, type, quantity, and position of labware on the Biomek i7 and peripheral instruments, as well as associated method details. SAMI EX scheduled paths and timing to shorten hardware movements, simulated the workflow to estimate time, and multitasked methods when the sample sizes are scaled up (Figure 1E). Steps such as plate transfers, cell imaging, sequential PBS washes, and the addition of trypsin and media were developed and managed through SAMI (Figure 1C). In contrast, Biomek 5 was used for more specific liquid handling steps, including cell density normalization and media backfilling (Figures 1C and 1G), with a dedicated deck layout (Figure 1F). The complete cell passaging workflow required about 32 min for one 96-well plate and 61 min for two plates when steps were overlapped (Figure 1E, Table 1).

**Table 1.** Time required for each automation pipeline or workflow.

|  | **1 plate** | **2 plates** | **4 plates** | **8 plates** |
| --- | --- | --- | --- | --- |
| **Mammalian cell culture passaging workflow.** | 32 min | 61 min | N/A | N/A |
| **Mammalian cell monoclonality screening pipeline.** | 4 min | 8 min | 16 min | 32 min |
| **Mammalian clonal line passaging and cherry-picking pipeline.** | 38 min | N/A | N/A | N/A |
| **Mammalian clonal cell number normalization for assay analysis and clone validation pipeline.** | 21 min | 39 min | N/A | N/A |
| **Mammalian clonal line cryopreservation pipeline.** | 5.5 min | 8.5 min | 12 min | 15 min |

N/A indicates 'not available' for the desired plate quantity due to limited deck capacity.

Reliable and reproducible cell number or confluency data from the parental plate is crucial for its passaging and for cell density normalization in daughter plates. At a high throughput scale, the CSI generates confluency data in less than 90 sec per 96-well plate (Figures 2B and 2C). This data, in CSV format, was automatically exported from CSI to the Biomek i7 for calculation of appropriate cell seeding volume. The Biomek then utilized a custom formula (Material and Methods 2.1) to determine the sample transfer volume from the parental (source) well to the daughter (destination) well. This study utilized HEK293T (an easier cell type requiring 0.025% of trypsin treatment), HEK293 (a sticker cell type requiring 0.25% of trypsin treatment), and HS-SY-II (a heterogenous cell type requiring 0.25% of trypsin treatment) cells with multiple biological replicates to demonstrate the workflow consistency. After HEK293 cell passaging and density normalization, most wells of the quadruplicate daughter 96-well plates showed comparable confluency within three standard deviations, indicating 99.7% of samples were effectively normalized with an average coefficient of variation (CV) of 16%, confirmed by CellTiter-Glo (CTG) viability assay and CSI confluency imaging (Figure 2A). To evaluate the workflow under extreme conditions and test normalization efficiency, an 8-point, 2-fold serial dilution was performed to establish a cell density gradient of HEK293 and 293T cells ranging from 100% to 0.8% confluency in a 96-well format, with 12 replicate wells per dilution. Due to the 128-fold difference in cell confluency between the top and bottom rows after cell seeding, prolonged incubation allowed cells in the lower density wells to expand, but skewed the median density toward higher confluency before normalization (Figure 2B).

Meanwhile, higher density wells in upper rows became overcrowded and compact, leading to a challenging condition to measure and normalize cell density. Therefore, one round of cell passaging and normalization can only reduce CV values by around 2 folds, from an average of 60% in the parental plates to 30% in the daughter plates for HEK293 cells (Figure 2B), and from 80% to 40% for 293T cells (Figure 2C). More importantly, the normalization procedure yielded a much narrower interquartile range, with the spread between Q1 and Q3 reduced from 70% to 5%, and a more symmetric confluency distribution across wells (Figures 2B and 2C).

### Arrayed mammalian cell monoclonality screening workflow

The high throughput arrayed clonal line establishment workflow is suitable for most mammalian cell types. To validate the workflow, HEK293T cells served as an easier cell model with high monoclonality, and HS-SY-II cells with a HiBiT tag KI represented a more difficult, heterogeneous cell line to grow clonally. The workflow consists of four key steps, including screening for cell monoclonality, passaging and cherry-picking colonies, normalizing cell numbers for assay analysis and clone validation, and cryopreserving the clonal lines.

Firstly, SAMI EX controlled the pipeline design (Figure 3A), method creation (Figure 3B), and coordinated with CSI for monoclonality screening of two 384-well plates. Additionally, SAMI scheduled and executed the pipeline in 8 min, processing two plates independently without parallel tasking (Figure 3C, Table 1). The labware setup report which illustrated the initial deck layout for the cell screening is shown in Figure 3D. To generate a monoclonality report (Figure 4), the same screening pipeline had to be repeated for one to two weeks in order to track cells from a single seeded state to expanded clonal lines. The well confluency data displayed in both numeric and heat map formats allows easy selection of wells containing cells above a certain confluency threshold (Figure 4A). Additionally, cell confluency images presented in thumbnail format enable the visual identification of candidate wells with single colony morphology (Figure 4B). The monoclonality result can be validated via time stamped images displayed chronologically (Figure 4D). Taken together, a full-well image for well B20 as an example of HEK293T cell was captured 10 days post seeding (Figure 4C). While the default algorithm offered a fixed set of 81 fields per well for cell migration and proliferation tracking via time stamped images, a user definable field could be designated to ensure capture of a single cell from initial seeding through colony formation (Figure 4E). The CSI can further analyze serial cell confluency data and create clonal growth curves to identify the most robust clones with exponential proliferation (Figure 4F).

After monoclonality assessment, the fast growing clonal lines in parental 384-well plates required to be passaged, cherry-picked, and reformatted into a daughter 96-well plate (Figure 6A), This study utilized HS-SY-II HiBiT reporter CRISPR KI clonal candidate cells as a model. SAMI EX continued to govern most of the pipeline design (Figure 5A) and method development (Figure 5B), except for the PBS replacement and cell suspension mixing steps. These steps were performed using a 384-channel pipette head and their deck configuration (Figure 5E) and methods (Figure 5F) were implemented using the Biomek 5 software, marked by the asterisk symbol in Figure 5B. Subsequently, flexible 8-channel pipette heads were utilized to cherry-pick individual clones and reformat them to a new plate. SAMI scheduled and completed the entire pipeline from a 384-well plate to a 96-well plate in 38 min (Figure 5C, Table 1). Its labware setup report summarized the initial deck layout and labware details (Figure 5D). When cells reached high confluency (Figure 6A), the daughter 96-well plate required an additional cell culture passaging workflow (Figure 1) to normalize cell density in a granddaughter plate (Figure 6A) for CTF and HiBiT assay analysis (Figure 6B). The normalized HiBiT/CTF values helped identify clonal candidates which had a higher likelihood of site specific KI, so the time and cost of downstream genomic DNA sequencing analysis can be reduced (Figure 6B). Meanwhile, the remaining cell suspension from the passaged daughter 96-well plate can be cryopreserved in a new 96-well plate for long-term storage (Figure 7). Runtimes for the cell cryopreservation pipeline were around 5.5 min for one 96-well plate and 12 min for 4 plates with overlapping procedures (Figure 7C, Table 1). The mammalian clonal line establishment workflow requires a total of 69, 132, 220, and 355 min to process one, two, four, and eight single cell seeded 384-well plates, respectively (Table 1). The arrayed mammalian cell monoclonality screening workflow enriched an average of 92 HS-SY-II reporter monoclones in a 96-well daughter plate, derived from 383 single cell seeded wells and one control well for cell focusing (100 cells seeded in the A1 position) (Figures 4B and 6A) in each 384-well parental plate. On average, the 96-well daughter plate yielded 35 clonal hits, determined from normalized HiBiT/CTF values, for downstream knock-in sequence analysis by genomic DNA sequencing (data not shown).

## Discussion

For most *in vitro* cancer cell models, high throughput arrayed cell culture automation remains the primary bottleneck limiting the duration of arrayed phenotypic screening using CRISPR gRNA, shRNA, or drug libraries to one week or less. To extend the screening timeframe, the present study focused on developing a standardized operating workflow to not only passage cells in the source 96-well plate format but also normalize cell density in the destination plate cost-effectively and consistently. Theoretically, this workflow is suitable for projects involving cell panel creation and maintenance. Furthermore, this workflow was optimized for clonal line screening and establishment, which is typically labor intensive, costly, and time consuming yet particularly valuable for CRISPR/Cas gene editing.

This study utilized a Biomek i7 Hybrid automated workstation to streamline two different cell culture workflows. For the hardware, this Biomek platform integrated a dual-pod liquid handler with peripheral instruments to maximize functionality and efficiency of automation. Accurate cell seeding number is the critical step for cell passaging, which commonly relies on the calculation of either cell density ratios or the exact cell counts. While other automated approaches, such as the Vi-CELL BLU counter (Beckman Coulter) for viability measurement and the Celigo image cytometer (Revvity) for cell density detection, are widely used, this study chose the CloneSelect Imager (CSI) for cell confluency analysis due to its rapid scanning capabilities. The CSI can accomplish a 96-well plate scan in 90 sec and a 384-well plate in 100 sec, which are crucial for high throughput automation. For the software, the Biomek 5 method utilized the built-in formula (Material and Methods 2.1) that processed auto-exported confluency data from CSI to calculate sample transfer volumes for cell density normalization in the destination plate. To normalize a 96-well plate, the passaging ratios were limited between 1:1 and 1:10. This range was selected to minimize the impact of outlying values and to maintain pipetting accuracy during cell suspension transfer. To enhance cell dissociation efficiency, the PBS wash step can be performed before trypsinization to remove residual media. Although the tilting-plate technology allows liquid removal and suspension mixing using flexible 8-channel pipettes, its application is limited at lower throughputs. Therefore, the BioTek 405 LSHTV washer was utilized in this study due to its speed and effectiveness in liquid aspiration; however, it was not used for liquid dispensing because of its tendency to disrupt the cell monolayer. To reduce costs and minimize contamination, the multichannel wash station (MC wash, Figure 1A) was implemented to rinse 96-well pipette tips with molecular-grade water, which enables tip re-usage. Cell dissociation during trypsinization was facilitated by the BioShake 3000-T ELM microplate shaker, which maintained a 37°C incubation temperature and dissociated the remaining adherent cells by vortexing. Complete cell dissociation after trypsin neutralization was achieved by vigorously pipetting at five distinct positions within each well, which was the most effective and efficient approach. For the software, SAMI EX and Biomek 5 provided user-friendly interfaces for method development and workflow management, planning, execution, and tracking. Experimental processes were tightly controlled from the entire workflow down to individual methods in spatial and temporal dimensions. SAMI EX, a dynamic scheduler, can efficiently estimate time and allocate resources for each method to facilitate multitasking. The workflow, pipeline, and method can be simulated in 2D or 3D, which enables visualization of software scheduling, hardware movement, and experiment timeline. The *in silico* demos are crucial to validate the hardware functionality, optimize protocols, and identify potential errors before physical execution. Additionally, SAMI’s error detection and recovery mechanisms, including proactive intervention, real-time user notification, and predefined corrective action, can significantly enhance the automation reliability and system uptime.

The arrayed cell passaging and density normalization workflow in this study reduced the interquartile range of cell density from 70% to 5% and the CV% from 60% and 80% in the parental plates to 30% and 40% in the daughter plates, respectively (Figures 2B and 2C). The residual high CV after normalization resulted from the initial 128-fold difference in cell confluency between the top and bottom rows after cell seeding, because the normalization workflow was designed to correct density variations within a 10-fold range (Material and Methods 2.1). Wells seeded at less than 3% confluency were unable to reach the target density even when the entire cell suspension was transferred, whereas over-confluent wells were likely underestimated in cell numbers based on confluency measurements, resulting in excess cells being allocated to the daughter wells during normalization. In comparison, direct cell seeding from a single source suspension using a dispenser, which is considered the most accurate approach to equalize cell number across wells, still resulted in an average CV of 7%, measured by cell viability (CTG assay) and cell density (CSI imaging) (data not shown). The Biomek i7 workflow, yielding an average CV of 16% after cell normalization (Figure 2A), was therefore considered acceptable because the pipetting and sampling errors from a 96-channel pipette head are higher than those from a liquid dispenser. Additionally, the similar CV values obtained from CTG and CSI indicated that cell confluency measured by CSI imaging is consistent in precision and reliability with viability measurements (Figure 2A). On the other hand, the arrayed mammalian cell monoclonality screening workflow, which leverages the high throughput imaging capability of the CSI and the cherry-picking function of the Biomek i7 liquid handler, together with the arrayed cell passaging and density normalization workflow, substantially enhanced clonal cell line establishment under scalable conditions (Table 1), while reducing labor, time, and errors caused by human intervention. Taken together, this study employed the Biomek i7 Hybrid automated workstation and integrated instruments to automate high throughput arrayed mammalian cell line cultivation workflows, which directly address unmet needs for arrayed library screening and clonal line enrichment post-mutagenesis.

## Acknowledgement

CH. and CY were supported by the Sanford Burnham Prebys (SBP) NCI Cancer Center Support Grant P30 CA030199. Research reported in this publication was supported by the SBP Functional Genomics Core through NIH Shared Instrumentation Grant S10 OD036254. Additional support was provided through NIH grants R01 AR078559 and R01 AG071861. We would like to acknowledge the Beckman Coulter team, including Brandon K. Corbin, Kenzo Maetani, Michael D. Moran, Eugene B. Tupas, and Liz Chladny, for their hardware support.

## Author contributions

CH, AB, AJD, MJ, and PDA conceptualized and designed the studies. DL, AVG, ZKW, JAY, and CY participated in methodology development and contributed to the automation workflow establishment. CY, DL, and CH performed the experiments and analyzed data. CH, CY, and AB wrote and edited the manuscript.

